# Morphine-induced mechanical hypersensitivity in mice requires δ receptors, β-arrestin2 and c-Src activity

**DOI:** 10.1101/2022.12.16.520707

**Authors:** Samuel Singleton, Tim G. Hales

## Abstract

**Background:** Morphine diminishes acute pain, but long-term use is compromised by tolerance and hyperalgesia. Studies implicate δ receptors, β-arrestin2 and Src kinase in tolerance. We examined whether these proteins are also involved in morphine-induced hypersensitivity (MIH). A common pathway for tolerance and hypersensitivity may provide a single target to guide improved analgesic approaches.

**Methods:** We examined mechanical sensitivity using automated von Frey in wild type (WT) and transgenic male and female C57Bl/6 mice before and after hind paw inflammation by complete Freund’s adjuvant (CFA). We explored the expression of opioid genes in the spinal cord using quantitative RT-PCR.

**Results:** CFA-evoked hypersensitivity ceased on day 7 in WT mice but persisted in μ^-/-^ mice. Recovery was delayed until day 13 in δ^-/-^ mice. Restoration to basal sensitivity in WT mice occurred with increased δ expression. By contrast, κ expression was reduced, while μ remained unchanged. Daily morphine reduced hypersensitivity in WT mice on day 3 compared to controls, however hypersensitivity recurred on day 9 and beyond. By contrast, WT mice had no recurrence of hypersensitivity in the absence of daily morphine. We used β-arrestin2^-/-^, δ^-/-^ and Src inhibition by dasatinib in WT mice to establish whether these approaches, which diminish tolerance, also attenuate MIH. While none of these approaches affected CFA-evoked inflammation or acute hypersensitivity, all caused sustained morphine anti-hypersensitivity, abolishing MIH.

**Conclusions:** Like morphine tolerance, MIH in this model requires δ receptors, β-arrestin2 and Src activity. Our findings suggest that MIH is caused by a tolerance-induced reduction in endogenous opioid signalling.

## Introduction

Morphine remains a first line treatment for severe acute pain 200 years after its first commercial production (Brook et al., 2017). However, its long-term use is compromised by tolerance and morphine induced hypersensitivity (MIH) (Colvin et al., 2019). A single approach that could diminish both effects would improve opioid analgesia in the context of persistent pain, but it remains unclear whether these phenomena are interdependent. While some studies suggest that morphine antinociceptive tolerance leads to MIH (Mao et al., 1995; Corder et al., 2017), others identify unrelated mechanisms (Juni et al., 2006; Roeckel et al., 2017).

Activation of μ receptors is required for morphine antinociception (Matthes et al., 1996). The development of morphine tolerance is influenced by the number of μ receptors as evidenced by the finding that μ^+/-^ mice, which express 50% fewer μ receptors than μ^+/+^ mice (Matthes et al., 1996), develop more rapid and profound morphine tolerance (Sora et al., 2001; Bull et al., 2017a). Morphine tolerance is also influenced by δ receptors. Tolerance is reduced by approaches that diminish δ receptor expression, function (Zhu et al., 1999; Nitsche et al., 2002) and association with μ receptors (Xie et al., 2009; He et al., 2011). The latter implies that δ receptors contribute to morphine tolerance by forming μ-δ heteromers (Gomes et al., 2000). Interestingly, the co-expression of μ and δ receptors leads to constitutive recruitment of β-arrestin2 which may prime morphine tolerance (Rozenfeld and Devi, 2007; Bull et al., 2017b).

When activated by agonists, μ receptors are targeted by GRK-mediated phosphorylation which stimulates recruitment of β-arrestin2, promoting receptor internalisation and activation of several kinases including c-Src (Walwyn et al., 2007; Williams et al., 2013). Constitutive β-arrestin2 knockout or modification of μ receptors to prevent GRK-mediated β-arrestin2 recruitment reduces morphine tolerance in mice as does c-Src inhibition (Bohn et al., 2000; Bull et al., 2017a; Kliewer et al., 2019). While β-arrestin2 and c-Src participate in morphine tolerance their involvement in MIH is unknown (Colvin et al., 2019).

Experimental models of persistent inflammation reveal a role of opioid receptors in regulating hypersensitivity under inflammatory conditions (Gaveriaux-Ruff et al., 2008; Luo et al., 2008; Obara et al., 2009; Corder et al., 2013). Both μ and δ receptors participate in spontaneous recovery from mechanical hypersensitivity in mice administered complete Freund’s adjuvant (CFA) (Gaveriaux-Ruff et al., 2008; Gaveriaux-Ruff et al., 2011; Corder et al., 2013; Walwyn et al., 2016). Paw injection of CFA causes mechanical hypersensitivity that recovers in mice within a week despite persistent inflammation. Hypersensitivity can be restored by injection of naltrexone, consistent with the idea that recovery reflects an upregulation of endogenous opioid signalling (Corder et al., 2013).

We used CFA-induced mechanical hypersensitivity to test the hypothesis that δ receptors, β-arrestin2 and c-Src kinase are required for MIH. We used strategies that accelerate (heterozygous µ^+/-^ mice) and attenuate (β-arrestin2^-/-^ and δ^-/-^ mice or c-Src inhibition in WT mice) tolerance to investigate its relationship to MIH in this model. We also determined the impact of persistent inflammation on the expression of transcripts encoding opioid peptide precursors, opioid receptors, β-arrestin2 and C-terminal Src kinase (CSK) in the spinal cord of WT mice to determine their potential contribution to inflammatory hypersensitivity.

## Methods

### Animals

Male and female wild type, µ^+/-^, µ^-/-^, δ^-/-^ or β-arrestin2^-/-^ mice aged eight- to 20-weeks old weighing between 18 g – 29 g when assigned to experimental treatment protocols were generated using a previously described breeding regime (Bull et al., 2017a; Bull et al., 2017b). All mice were maintained on the C57BL6/J background and housed in social groups comprised of same-sex littermates under standard conditions in a 12-hour dark-light cycle with free access to food and water. All experiments were performed on mice of both sexes born from multiple litters, with approximately equal numbers coming from each cage and sex where possible. At the end of the experiment, mice were culled via cervical dislocation and genotypes confirmed using the automated genotyping service provided by Transnetyx (USA). All studies were reviewed and approved by the Ethical Committee in the University of Dundee’s Medical School Resource Unit (Dundee, United Kingdom) and comply with United Kingdom Home Office regulations. Additionally, the use and reporting of animals within this manuscript complies with the ARRIVE guidelines version 2.0 (Percie du Sert et al., 2020).

### Inflammation

Mice were unilaterally injected (intraplantar) with 10 µL of undiluted (1 mg/mL) CFA (Sigma, UK) into one hind paw using a 100 µL syringe and 29-gauge needle. The vertical distance from the first walking pad to the ventral surface of each hind paw was measured using FiJi (ImageJ v1.51n) with individually scaled images. Inflammation was monitored by subtracting the width at baseline from the observed width of each paw on each day (Δ footpad width). Recovery from inflammation was measured by subtracting the width measured on the final day from the width measured 1 day after injection with CFA (ΔΔ footpad width). Data are presented as the change in absolute paw widths (in mm) ± SD.

### Mechanical sensitivity

Mice were individually housed in plexiglass chambers on an elevated mesh platform. Mechanical sensitivities of each hind paw were measured using a dynamic plantar aesthesiometer (Ugo Basile, Italy) set to deliver an increasing force at a linear rate of 2.5 g/s. Responses of each hind paw were measured three times on each day and the average for each paw calculated as the withdrawal threshold. Baseline responses are the mechanical sensitivities measured immediately prior to injection with CFA and represent the final assessment after three consecutive days of testing to habituate the mice to the testing chamber. Mechanical sensitivities were reassessed 24 hours following injection with CFA and once every 48 hours thereafter. Mice were left to habituate to the testing chamber for 30 minutes prior to each assessment. All measurements were performed by the same individual blinded to treatment and/or genotype. Data represented in graphs are expressed as absolute withdrawal thresholds (in g) ± SEM. Mechanical sensitivity data presented in the corresponding Tables were additionally converted to the percentage change from baseline sensitivity using the calculation 100*(observed threshold – baseline threshold)/baseline threshold.

### Drug preparation and procurement

Drugs for *in vivo* experiments were prepared immediately before use in sterile conditions and filtered using a 0.2 µm syringe. Morphine sulfate (Sigma, UK) was prepared as a 25 mg/mL stock solution in sterile water and reconstituted in 0.9% saline to a final concentration of 2 mg/mL. Dasatinib (Bristol-Myers Squibb, USA) was prepared as a 50 mg/mL stock solution in DMSO and reconstituted in 0.9% saline to a final concentration of 1 mg/mL and supplemented with 2% Kolliphor EL (BASF Pharma, USA). Morphine or matched saline injections were administered subcutaneously into the scruff of the neck while injections with dasatinib or matched vehicle (0.9% saline with 2% v/v Kolliphor EL) were administered via the intraperitoneal route. All injections were administered 30 minutes prior to assessing mechanical sensitivity. In the case of morphine and dasatinib co-administration, dasatinib was injected 30 minutes prior to morphine each day. Mice were randomly assigned to receive experimental or control treatments prior to each experiment.

### Assessing hypersensitivity

Spontaneous recovery from CFA-evoked mechanical hypersensitivity was assessed in the absence of vehicle injections. In contrast, to assess morphine anti-hypersensitivity and morphine-induced hypersensitivity (MIH), mice received once-daily injections with morphine (3 mg/Kg or 10 mg/Kg) or saline (vehicle) starting two days after injection with CFA to establish and confirm the development of mechanical hypersensitivity prior to morphine treatment. Morphine anti-hypersensitivity and MIH were assessed within each genotype using absolute withdrawal thresholds, or, compared between genotypes using mechanical sensitivities that were converted to a percentage reduction from baseline sensitivities to compensate for altered baseline mechanical sensitivities. Furthermore, the change in mechanical sensitivity on day 15 compared to baseline sensitivity was additionally used to compare MIH between mice receiving repeated injections with morphine to those receiving vehicle.

### Western blots

Lumbar (L1-L5) spinal cord tissue collected from mice post-mortem was mechanically homogenised followed by sonication on ice. Tissue lysis was performed in ice cold RIPA buffer (Abcam, UK) supplemented with a protease inhibitor cocktail (Roche, UK) and Na^+^ orthovanadate (1 mM). Samples were centrifuged for 20 minutes (12,000 rpm at 4°C) and the protein concentration of each supernatant was determined using the Pierce BCA assay (Thermo Fisher, UK). Proteins were denatured by 2-mercaptoethanol at 95°C and separated using SDS-PAGE in MOPS running buffer on a NuPAGE Bis-Tris 4-12% gel (ThermoFisher, UK). Separated proteins were transferred to nitrocellulose membranes (GE healthcare, UK) and probed with rabbit anti-c-Src primary antibodies raised against c-Src in its non-phosphorylated (1:2,000; Cell Signaling, UK) or phosphorylated form (1:1,000; Cell Signaling), or GAPDH (1:2,000; Abcam, UK). Protein bands were detected using a horseradish peroxidase conjugated goat anti-rabbit secondary antibody (1:5,000 - 10,000; Abcam) and visualised by enhanced chemiluminescence (GE Healthcare, UK) on AGFA medical X-ray film (blue). Densitometric quantification of c-Src in either its phosphorylated or non-phosphorylated form was determined using Fiji ImageJ (v1.51n) and expressed relative to GAPDH for each membrane.

### Quantitative PCR

Lumbar (L1-L5) spinal cord tissue was rapidly collected from mice post-mortem in RNase free conditions and snap frozen in liquid N_2_. Samples were thawed in TRIzol (ThermoFisher, UK) and sonicated prior to RNA extraction by chloroform and isopropanol. Precipitated RNA was resuspended in nuclease free water and 1 µg total RNA was reverse transcribed into cDNA using SuperScript™ II (Invitrogen, UK) as per manufacturer’s instructions. RT-qPCR was performed on a QuantStudio™ 7 Flex System (Applied Biosystems, UK) in a total reaction volume of 20 µL comprised of 10 µL TaqMan universal PCR mastermix-II (ThermoFisher, UK), 6.5 µL nuclease free ddH2O (Ambicon, UK), 2.5 µL cDNA and 1 µL TaqMan gene expression array primers (ThermoFisher, UK). mRNA expression was determined after 40 cycles by the Δ cycle threshold (CT) method (2^--ΔΔCT^) using GAPDH as the reference gene. Relative expression data for each target mRNA are presented as the fold change over control expression.

TaqMan array predesigned gene expression primers were used to assess *Oprm1* (µ opioid receptor; Mm01188089_m1), *Oprd1* (δ opioid receptor; Mm01180757_m1), *Oprk1* (κ opioid receptor; Mm01230885_m1), *Pomc* (proopiomelanocortin; Mm00435874_m1), *Penk* (proenkephalin; Mm01212875_m1), *Pdyn* (prodynorphin; Mm00457573_m1), *Arrb2* (β-arrestin2; Mm00520666_g1), *Csk* (c-terminal Src kinase; Mm00432757_g1) or *Gapdh* (Glyceraldehyde 3-phosphate dehydrogenase; Mm99999915_g1) expression. All primers used a 5’ FAM reporter dye and 3’ nonfluorescent quencher.

### Statistics and data analysis

Data were imported into GraphPad Prism (v.5.0; USA) and IBM SPSS Statistics (v.27.0; USA) to construct graphs and perform statistical comparisons. Normality and homogeneity of variance for each dataset was confirmed prior to statistical analysis using the Shapiro-Wilk test and Levene’s box test, respectively. Spontaneous recovery from inflammatory hypersensitivity was assessed within each genotype using a one-way ANOVA with repeated measures on absolute withdrawal thresholds. Spontaneous recovery in transgenic mice was also compared to wild type mice using a two-way ANOVA with repeated measures on withdrawal thresholds that had been converted to a percentage of baseline sensitivities. Recoveries involving the daily administration of morphine and/or dasatinib were compared to vehicle treated mice within each genotype using a two-way ANOVA with repeated measures. Statistical comparisons between morphine and saline treated mice within each genotype were performed using the two-sample Student’s t-test, while comparisons between genotypes were performed using a two-way ANOVA. Multiple comparisons following detection of statistically significant F values were controlled for with Dunnett’s or Bonferroni *post hoc* corrections applied where appropriate. Group sizes specified in each Figure legend represent numbers of individual mice. Summary data are presented as mean ± SEM, or the mean ± SD where appropriate. Statistical comparisons were considered significant when p < 0.05.

## Results

### Inflammatory hypersensitivity is resolved through enhanced opioid signalling

We employed a model of persistent inflammation in male and female WT, μ^-/-^, δ^-/-^ and β-arrestin2^-/-^ mice by injecting each with 10 µL CFA (1 mg/mL) into one hind paw. Mechanical sensitivities were determined at baseline and reassessed for up to 15 days following injection with CFA in intervals of up to 48h (Figure 1A). Importantly, there was no change in the mechanical thresholds of the hind paw contralateral to the injection site nor were there any significant differences in recovery times between male and female mice in any genotype (data not shown). Therefore, data represent the mechanical sensitivities of the ipsilateral paw that have been pooled between male and female mice.

**Figure 1.**
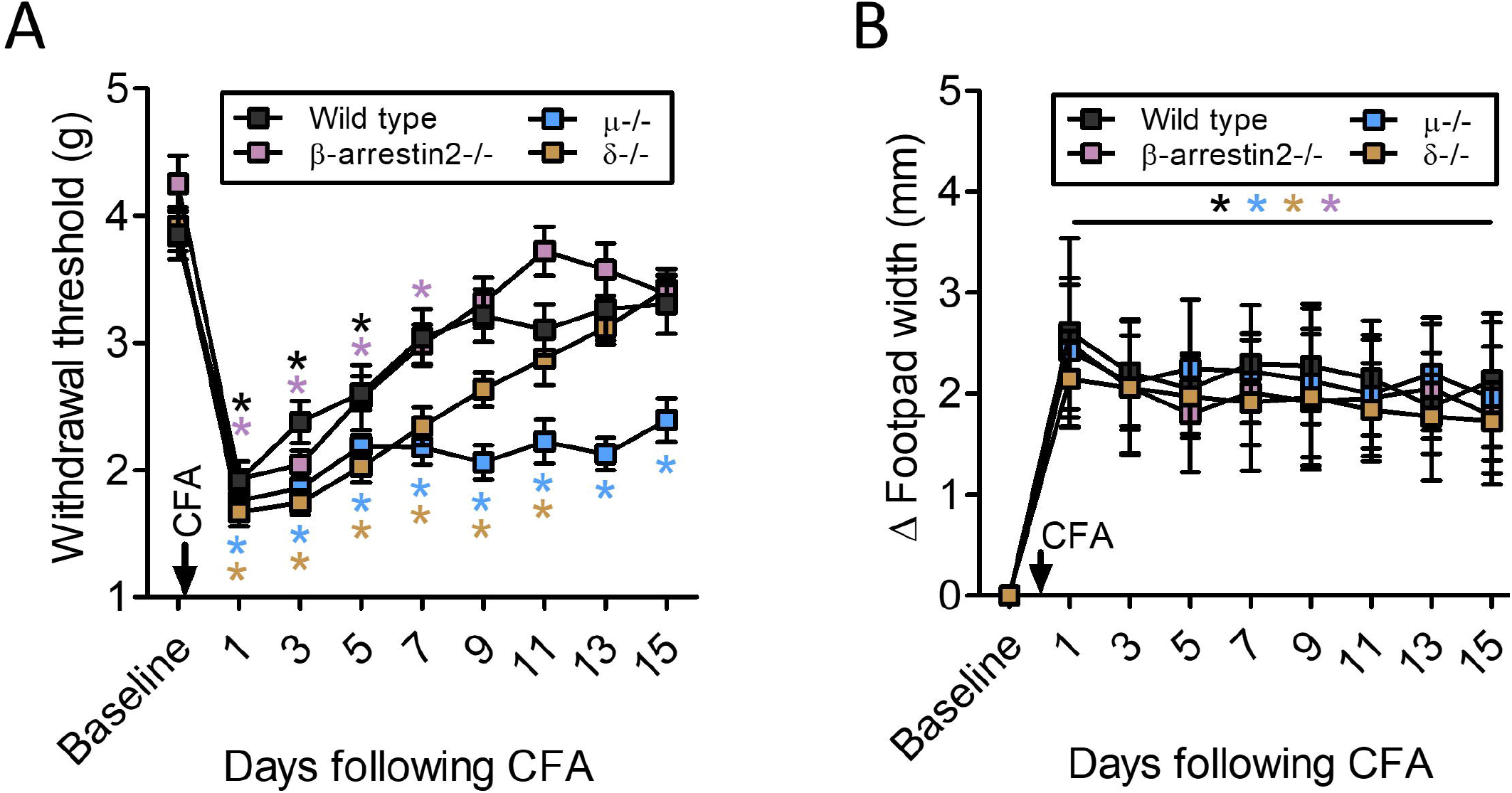
CFA causes transient mechanical hypersensitivity resolved through endogenous opioid signalling. (A) Development of CFA-evoked mechanical hypersensitivity in the ipsilateral hind paws of WT, μ^-/-^, δ^-/-^ and β-arrestin2^-/-^ mice. Mechanical sensitivities are summarised in Table 1. A two-way ANOVA with repeated measures revealed that WT mice develop transient mechanical hypersensitivity that spontaneously recovers back to baseline sensitivity by day 7. In contrast, μ^-/-^ mice do not recover. Spontaneous recovery to basal withdrawal thresholds is delayed until day 13 in δ^-/-^ mice and until day 9 in β-arrestin2^-/-^ mice. (B) The change in hind paw width on each day following injection with CFA compared to baseline. A two-way ANOVA with repeated measures revealed a significant effect of time (F8,344 = 110.2; p < 0.0001) but no effect of genotype (F3,344 = 1.1; p = 0.37). Hind paw widths were determined by subtracting the dorsal-ventral distance, at baseline, from the distance measured on each day (Δ Footpad width). Data in (A) represents mean ± SEM while data in (B) represents mean ± S.D. n = 11 – 13 (WT), n = 10 (μ^-/-^), n = 11 (δ^-/-^) and n = 15 (β-arrestin2^-/-^). *p < 0.05 compared to baseline.

CFA caused the development of mechanical hypersensitivity in all mice as evidenced by a reduction in mechanical thresholds in the ipsilateral hind paw following injection (Figure 1A; Table 1). The development of mechanical hypersensitivity (mean percentage change ± SEM in all cases) was similar between WT (48 ± 4%), μ^-/-^ (54 ± 4%), δ^-/-^(57 ± 3%) and β-arrestin2^-/-^ (53 ± 3%) mice on day 1 following injection with CFA.

**Table 1:** Development of CFA-evoked mechanical hypersensitivity in WT, μ^-/-^, δ^-/-^ and β-arrestin2^-/-^ mice.

|  | Mechanical hypersensitivity (% reduction from baseline) |  |  |  |
| --- | --- | --- | --- | --- |
| Time after CFA (days) | Wild type | $\mu^{-/-}$ | $\delta^{-/-}$ | β-arrestin2$^{-/-}$ |
| Baseline | 0 | 0 | 0 | 0 |
| 1 | 48 ± 4* | 54 ± 4* | 57 ± 3* | 53 ± 4* |
| 3 | 36 ± 5* | 52 ± 3*<sup>#</sup> | 56 ± 2*<sup>#</sup> | 50 ± 3* |
| 5 | 29 ± 5* | 43 ± 3*<sup>#</sup> | 48 ± 3*<sup>#</sup> | 34 ± 10* |
| 7 | 17 ± 9 | 43 ± 4*<sup>#</sup> | 40 ± 4*<sup>#</sup> | 25 ± 8* |
| 9 | 14 ± 6 | 46 ± 3*<sup>#</sup> | 33 ± 2*<sup>#</sup> | 16 ± 9 |
| 11 | 17 ± 6 | 42 ± 5*<sup>#</sup> | 26 ± 6* | 7 ± 8 |
| 13 | 12 ± 9 | 45 ± 3*<sup>#</sup> | 19 ± 5 | 13 ± 6 |
| 15 | 12 ± 7 | 37 ± 4*<sup>#</sup> | 12 ± 5 | 16 ± 7 |

We assessed the duration of CFA-evoked mechanical hypersensitivity by monitoring the time taken to recover back to absolute baseline sensitivities within each genotype (Figure 1A) or across genotypes using mechanical thresholds that had been converted to a percentage decrease from baseline (Table 1). The latter approach allowed us to assess the contribution of endogenous opioid signalling to recovery from CFA-evoked mechanical hypersensitivity while accounting for potential differences in baseline mechanical sensitivities between genotypes. Hypersensitivity was transient in WT mice lasting up to 7 days whereas persistent mechanical hypersensitivity developed in μ^-/-^mice that lasted for all measurements following injection with CFA. In contrast, CFA-evoked mechanical hypersensitivity lasted for up to 13 days in δ^-/-^mice and for up to 9 days in β-arrestin2^-/-^ mice (Figure 1A).

A two-way ANOVA with repeated measures comparing recovery from mechanical hypersensitivity across genotypes (Table 1) detected a significant effect of time (F5,234 = 58.2; p < 0.001), genotype (F3,45 = 3.8; p < 0.001) and an interaction (F16,234 = 2.9; p < 0.001). Pairwise comparisons with the Bonferroni correction revealed that mechanical hypersensitivity in μ^-/-^ mice was significantly greater than that of WT mice from day 3 onwards. In contrast, mechanical hypersensitivity in δ^-/-^ mice was significantly greater than WT mice on day 3 to day 9. There was no significant difference in the mechanical sensitivities of β-arrestin2^-/-^ mice compared to WT mice at any point following injection with CFA (Table 1). Together, these data demonstrate that CFA-evoked mechanical hypersensitivity is transient in WT, β-arrestin2^-/-^and δ^-/-^ mice, although delayed in the latter. By contrast, there is no recovery from hypersensitivity in μ^-/-^ mice, consistent with a requirement for increased opioid signalling for restoration hypersensitivity to basal levels in WT mice.

### CFA-evoked hind paw inflammation is not affected in μ^-/-^, δ^-/-^ or β-arrestin2^-/-^ mice

We monitored the development of hind paw inflammation in each mouse to determine whether diminished spontaneous recovery in μ^-/-^ and δ^-/-^ mice (Figure 1A) was associated with altered inflammation (Figure 1B). Hind paw inflammation developed ipsilateral to the injection site in all mice following CFA while no change was observed in the contralateral hind paws (data not shown). Ipsilateral hind paw widths were increased (mean ± S.D in all cases) on day 1 by 2.6 ± 0.9 mm in WT mice, and by 2.4 ± 0.7 mm, 2.1 ± 0.5 mm and 2.5 ± 0.6 mm in μ^-/-^, δ^-/-^ and β-arrestin2^-/-^mice, respectively. On day 15, hind paw widths remained 2.1 ± 0.7 mm greater than baseline in WT mice, whereas they remained greater by 2.0 ± 0.7 mm, 1.7 ± 0.4 mm and 1.8 ± 0.7 mm in μ^-/-^, δ^-/-^ and β-arrestin2^-/-^ mice, respectively (Figure 1B). A two-way ANOVA with repeated measures comparing the change in hind paw widths, from baseline (Δ footpad width), across each genotype confirmed a significant increase on all days following injection with CFA (F8,344 = 110.2; p < 0.0001), although there was no effect of genotype (F3,344 = 1.1, p = 0.37). Furthermore, a one-way ANOVA comparing the change in footpad width on day 15 compared to day 1 (a measure of resolution) in all mice, similarly revealed no effect of genotype (F3,43 = 0.72; p = 0.8, data not shown). Therefore, CFA injection into the hind paw causes robust localised inflammation that is not obviously affected by the absence of μ receptors, δ receptors or β-arrestin2. This implies that spontaneous recovery is not due to altered inflammation in any of the genotypes examined.

### Complete Freund’s adjuvant alters gene expression in the spinal cord of WT mice

Since spontaneous recovery from CFA-evoked mechanical hypersensitivity requires endogenous opioid receptor function (Figure 1A), in keeping with prior studies (Corder et al., 2013; Walwyn et al., 2016), we explored whether there was altered expression of relevant genes. We used quantitative PCR to examine transcripts encoding opioid peptide precursors, opioid receptors, β-arrestin2 and C-terminal Src kinase within the spinal cord of WT mice (Figure 2). The latter regulates the activity of c-Src, a non-receptor tyrosine kinase that is implicated in both inflammatory hypersensitivity and morphine-evoked tolerance (Liu et al., 2008; Xu et al., 2012; Bull et al., 2017a; Chen et al., 2020).

**Figure 2.**
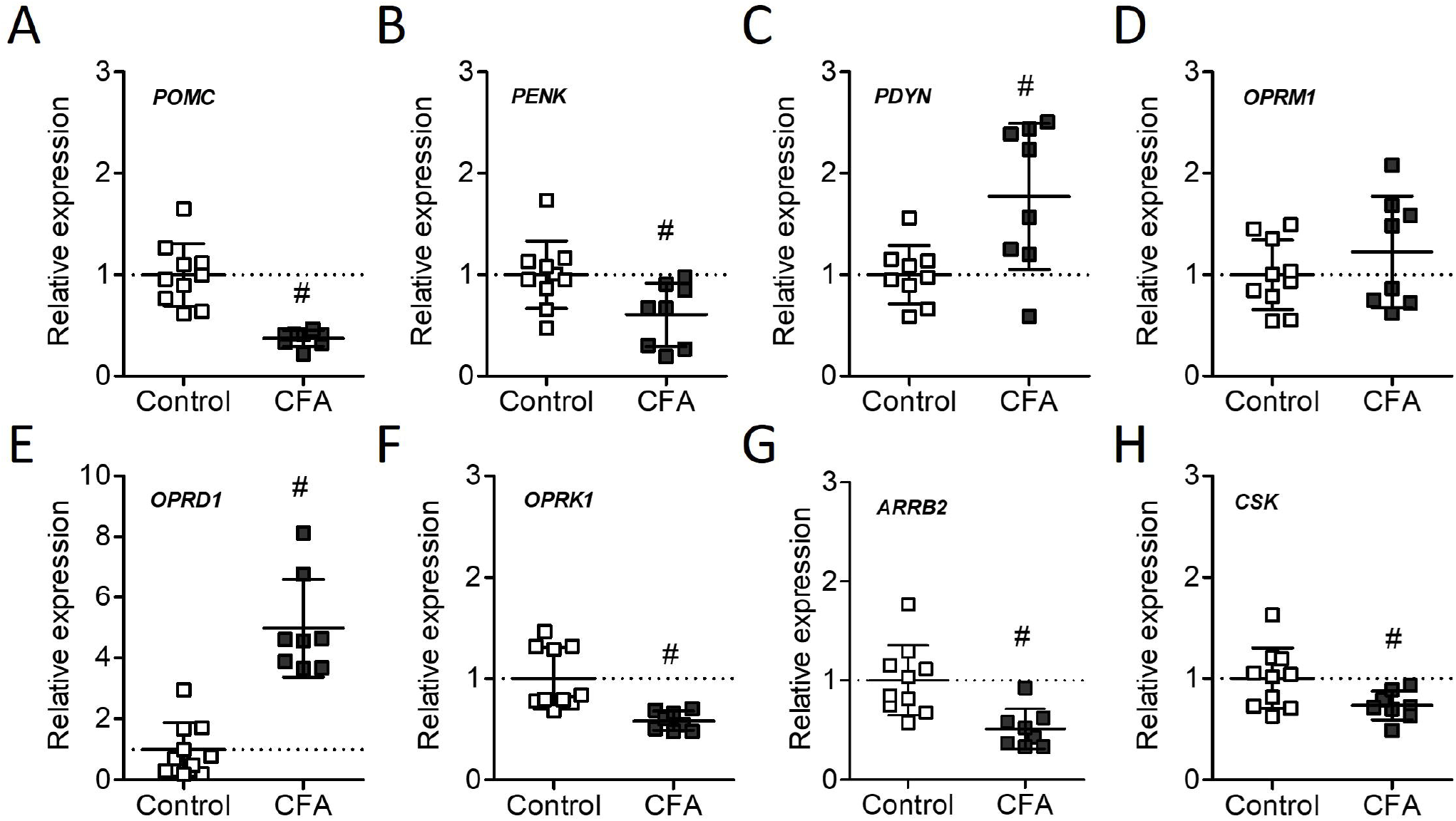
Altered opioid signalling in the spinal cord of WT mice injected with CFA. Expression of genes encoding opioid peptides proopiomelanocortin (A), proenkephalin (B), prodynorphin (C), opioid receptors µ (D), δ (E), κ (F), β-arrestin 2 (G) or c-terminal Src kinase (H) was quantified by qPCR in lumbar (L2-L5) spinal cord tissue harvested from WT mice on day 7 following a unilateral injection (intraplanter) with 10 μL CFA (1 mg/mL) or control. Gene expression was determined by the 2^-ΔΔCT^ method using GAPDH as the reference gene. Statistical comparisons were performed using the independent sample Student’s t-test (two-tailed). Expression of each gene is shown relative to control. Whisker plots represent mean ± SD. n = 10 (control), n = 8 (CFA). ^#^p < 0.05 compared to control.

Mechanical hypersensitivity is transient in WT mice lasting for 7 days (Figure 2A; Table 1), therefore, we injected CFA or an equal volume of saline (control) and harvested the lumbar spinal cords after 1 week. The mRNAs quantified were those transcribed from the *Pomc, Penk* and *Pdyn* (encoding proopiomelanocortin, proenkephalin and prodynorphin respectively), *Oprm1, Oprd1* and *Oprk1* (encoding the μ-, δ- and κ opioid receptor respectively), *Arrb2* (encoding β-arrestin2) and *Csk* (encoding C-terminal Src Kinase) genes.

CFA-evoked inflammation altered the expression of all genes tested that encode opioid peptide precursors (*Pomc, Penk* and *Pdyn*). The expression (mean ± S.D normalised to control expression in all cases) of *Pomc* (0.4 ± 0.02; p < 0.0001) and *Penk* mRNAs (0.6 ± 0.1; p = 0.021) were significantly reduced while the expression of *Pdyn* mRNA (1.8 ± 0.3; p = 0.009) was significantly enhanced in the spinal cord by injection with CFA (Figure 2A-C). The expression of *Oprm1* mRNA (1.2 ± 0.2; p = 0.31) was not significantly affected by CFA although *Oprd1* mRNA (5.0 ± 0.6; p < 0.0001) was significantly enhanced and *Oprk1* mRNA (0.6 ± 0.03; p = 0.002) was significantly reduced (Figure 2D-F). Interestingly, the expression of *Arrb2* mRNA (0.5 ± 0.1; p = 0.003) and *Csk* mRNA (0.7 ± 0.05; p = 0.034) were both significantly reduced in the spinal cords of CFA injected mice relative to control (Figure 2G-H). These data demonstrate that persistent inflammation and recovery from mechanical hypersensitivity are accompanied by altered expression of genes involved in opioid signalling mechanisms within the spinal cord.

### Reinstatement of mechanical hypersensitivity following repeated morphine injections in WT mice

We assessed the ability of an acute morphine injection (either 3 or 10 mg/Kg) to attenuate mechanical hypersensitivity in mice that had received a unilateral injection with CFA into one hind paw two days prior (Figure 3A). Morphine or an equal volume of saline was administered subcutaneously into the scruff of the neck and mechanical thresholds assessed after 30 minutes.

**Figure 3.**
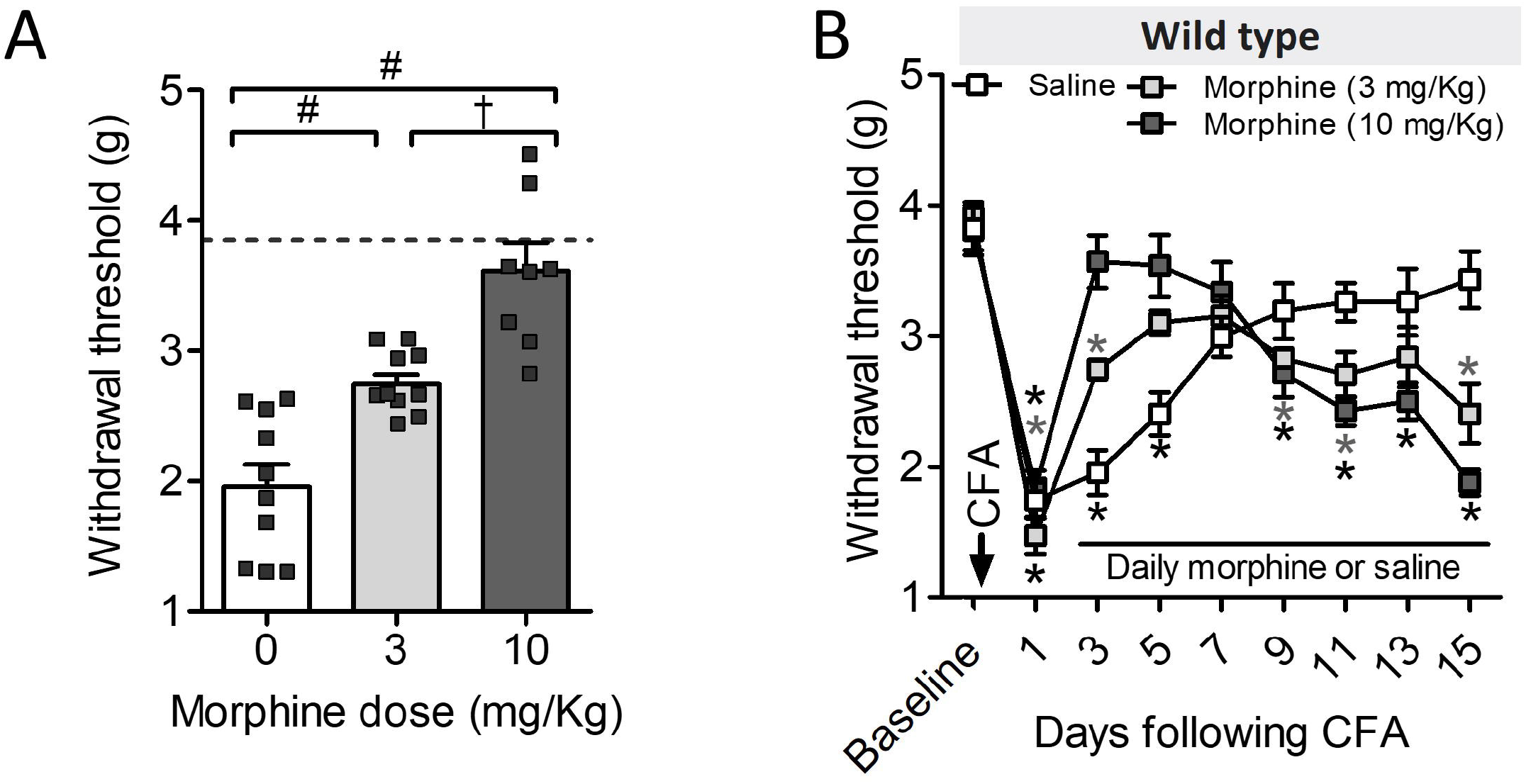
Morphine limits mechanical hypersensitivity but leads to its reinstatement following repeated injections in WT mice. (A) Acute morphine injection dose-dependently limits mechanical hypersensitivity caused by a 10 μL injection with CFA (1 mg/mL). A one-way ANOVA with Bonferroni pairwise comparisons revealed that morphine (3 mg/Kg) limited hypersensitivity compared to vehicle (saline) injections while morphine (10 mg/Kg) limited hypersensitivity compared to both vehicle and 3 mg/Kg morphine (F2,25 = 26.7, p < 0.0001). (B) Acute morphine anti-hypersensitivity transitions to persistent mechanical hypersensitivity following repeated injections. A two-way ANVOA with Bonferroni pairwise comparisons revealed that morphine initially limited hypersensitivity compared to vehicle on days 3 and 5 following injection with CFA. However, hypersensitivity compared to baseline recurred by day 9 after repeated morphine injections whereas baseline mechanical sensitivity was sustained following recovery in mice receiving vehicle. Statistical comparisons between treatment groups on each day are summarised in Table 2. Data represent mean ± SEM. (A) n = 10 (saline), n = 10 (3 mg/Kg) and n = 8 (10 mg/Kg). (B) n = 10 (saline) and n = 10 (morphine 3 mg/Kg). *p < 0.05 compared to baseline, ^#^p < 0.05 compared to saline, ^†^p < 0.05 dose effect.

Morphine dose-dependently attenuated mechanical hypersensitivity restoring baseline withdrawal thresholds completely following injection with 10 mg/Kg (Figure 3A). Withdrawal thresholds were increased by 40 ± 4% (3 mg/Kg) and 82 ± 10% (10 mg/Kg) relative to saline injected mice.

**Table 2:**
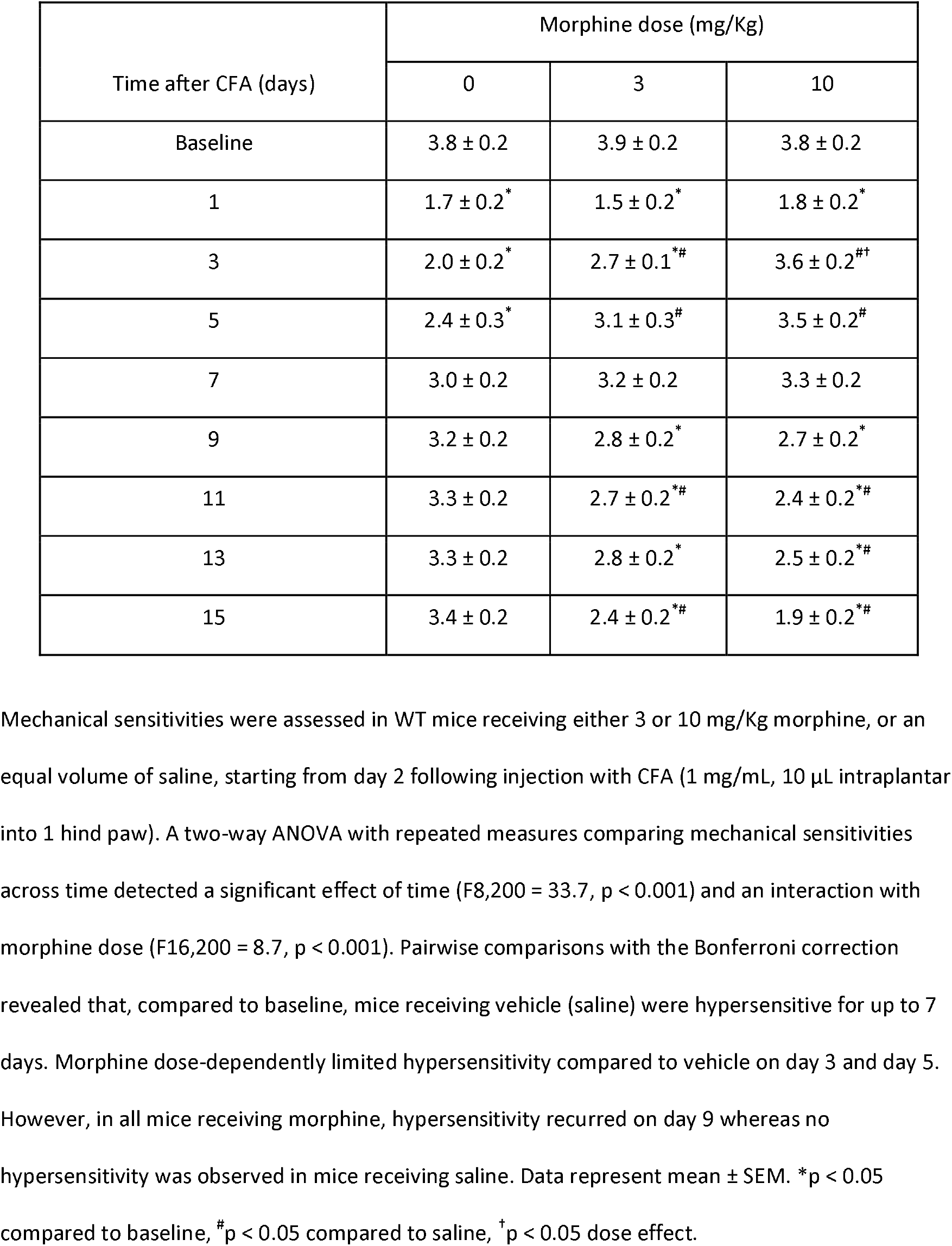
Morphine anti-hypersensitivity and morphine induced hypersensitivity in WT mice.

| Time after CFA (days) | Morphine dose (mg/Kg) |  |  |
| --- | --- | --- | --- |
|  | 0 | 3 | 10 |
| Baseline | 3.8 ± 0.2 | 3.9 ± 0.2 | 3.8 ± 0.2 |
| 1 | 1.7 ± 0.2<sup>*</sup> | 1.5 ± 0.2<sup>*</sup> | 1.8 ± 0.2<sup>*</sup> |
| 3 | 2.0 ± 0.2<sup>*</sup> | 2.7 ± 0.1<sup>*#</sup> | 3.6 ± 0.2<sup>#†</sup> |
| 5 | 2.4 ± 0.3<sup>*</sup> | 3.1 ± 0.3<sup>#</sup> | 3.5 ± 0.2<sup>#</sup> |
| 7 | 3.0 ± 0.2 | 3.2 ± 0.2 | 3.3 ± 0.2 |
| 9 | 3.2 ± 0.2 | 2.8 ± 0.2<sup>*</sup> | 2.7 ± 0.2<sup>*</sup> |
| 11 | 3.3 ± 0.2 | 2.7 ± 0.2<sup>*#</sup> | 2.4 ± 0.2<sup>*#</sup> |
| 13 | 3.3 ± 0.2 | 2.8 ± 0.2<sup>*</sup> | 2.5 ± 0.2<sup>*#</sup> |
| 15 | 3.4 ± 0.2 | 2.4 ± 0.2<sup>*#</sup> | 1.9 ± 0.2<sup>*#</sup> |

Mechanical sensitivities were assessed in WT mice receiving either 3 or 10 mg/Kg morphine, or an equal volume of saline, starting from day 2 following injection with CFA (1 mg/mL, 10 µL intraplantar into 1 hind paw). A two-way ANOVA with repeated measures comparing mechanical sensitivities across time detected a significant effect of time ( $F_{8,200} = 33.7$ , $p < 0.001$ ) and an interaction with morphine dose ( $F_{16,200} = 8.7$ , $p < 0.001$ ). Pairwise comparisons with the Bonferroni correction revealed that, compared to baseline, mice receiving vehicle (saline) were hypersensitive for up to 7 days. Morphine dose-dependently limited hypersensitivity compared to vehicle on day 3 and day 5. However, in all mice receiving morphine, hypersensitivity recurred on day 9 whereas no hypersensitivity was observed in mice receiving saline. Data represent mean ± SEM. \* $p < 0.05$ compared to baseline, # $p < 0.05$ compared to saline, † $p < 0.05$ dose effect.

We next assessed whether morphine continued to attenuate mechanical hypersensitivity evoked by CFA in WT mice following repeated once-daily injections (Figure 3B). Morphine (either 3 or 10 mg/Kg) or an equal volume of saline was similarly administered two days following CFA injection, and mechanical thresholds assessed in 48h intervals, 30 minutes after each injection.

Morphine (3 mg/Kg) initially elevated mechanical thresholds on day 3 and day 5, although on day 9, after receiving morphine injections for 7 consecutive days, mechanical hypersensitivity had recurred as evidenced by reduced mechanical thresholds compared to baseline. Increasing the daily morphine dose led to enhanced anti-hypersensitivity on day 3 and day 5 although mechanical hypersensitivity similarly returned by day 9. In contrast, the mechanical thresholds of WT mice receiving vehicle (saline) were reduced from baseline on days 1, 3 and 5. Recovery back to baseline sensitivity was complete by day 7 and sustained until the final measurement on day 15 (Figure 3B; Table 2). When compared to saline injected mice, the mechanical thresholds of mice receiving morphine (3 mg/Kg) were reduced on day 11 and day 15, while the mechanical thresholds of mice receiving morphine (10 mg/Kg) were reduced from day 11. Furthermore, increasing the dose of morphine led to greater reinstatement of hypersensitivity on day 15 (Figure 3B; Table 2).

Together, these data demonstrate that, while morphine initially attenuates the development of mechanical hypersensitivity during persistent inflammation, repeated daily administration causes reinstatement of mechanical hypersensitivity not seen in mice receiving vehicle, i.e., morphine induced hypersensitivity (MIH).

### Contribution of μ and δ receptors to the modulation of hypersensitivity by morphine

We determined the contribution of μ and δ receptors to morphine anti-hypersensitivity and MIH during persistent inflammation by assessing the mechanical sensitivities of μ^-/-^ and δ^-/-^ mice following repeated injections with morphine (Figure 4 A-B). Morphine (3 mg/Kg) was similarly administered once daily, starting from day 2 following injection with CFA.

**Figure 4.**
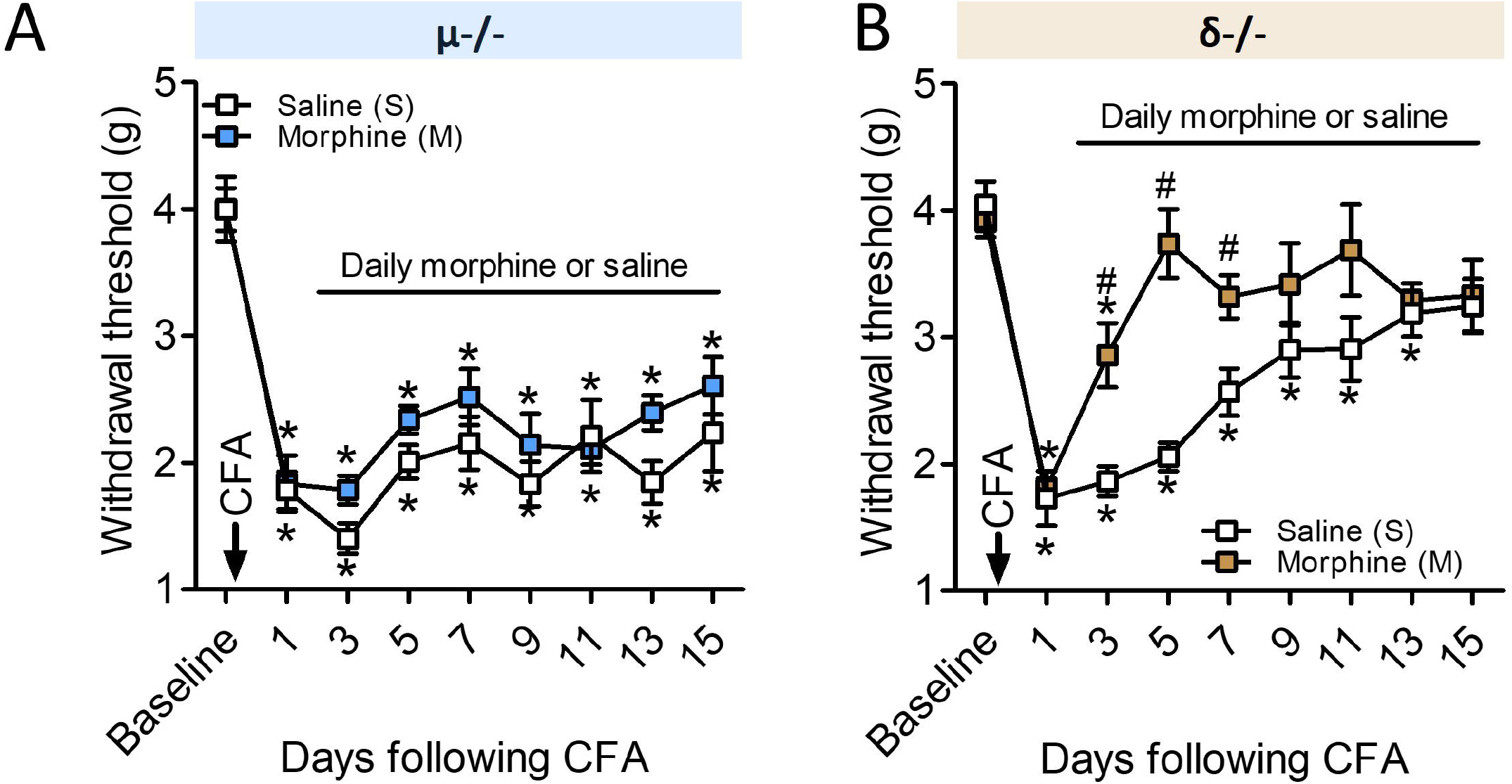
Morphine anti-hypersensitivity and transition to mechanical hypersensitivity in μ^-/-^ and δ^-/-^ mice. Mechanical sensitivities of (A) μ^-/-^ and (B) δ^-/-^ mice receiving once-daily injections with morphine (3 mg/Kg) or an equal volume of saline, starting from day 2 after an injection with 10 μL CFA (1 mg/mL) into one hind paw. (A) Morphine did not limit hypersensitivity or lead to mechanical hypersensitivity in μ^-/-^ mice (treatment: F1,18 = 1.2, p = 0.29; interaction: F2,36 = 0.6, p = 0.58). By contrast, morphine limited hypersensitivity in δ^-/-^ mice on days 3 – 7 compared to vehicle (B) although mechanical hypersensitivity did not recur despite repeated injections (treatment: F1,16 = 2.3, p = 0.14; interaction: F2,32 = 5.5, p = 0.01). Data represent mean ± SEM. n (saline then morphine) = 10 and 10 (µ^-/-^ mice), n = 11, 7 (δ^-/-^ mice). *p < 0.05 compared to baseline, ^#^p < 0.05 treatment effect.

Morphine had no effect on the mechanical sensitivities of μ^-/-^ mice (Figure 4A). The mechanical thresholds of μ^-/-^ mice on day 1 following injection with CFA (without morphine) were 1.7 ± 0.2 g while the mechanical thresholds on day 3 (after morphine) were 1.8 ± 0.1 g. These were similar to μ^-/-^ mice receiving vehicle on day 1 (1.8 ± 0.1 g) and day 3 (1.4 ± 0.1 g). A two-way ANOVA with repeated measures confirmed a significant effect of time following CFA injection (F8,144 = 33.9; p < 0.0001) although there was no significant effect of treatment (F1,18 = 2.4; p = 0.14), nor was there an interaction (F8,144 = 0.8; p = 0.58). Pairwise comparisons for the significant main effect of time with the Bonferroni correction revealed that, similar to spontaneous recovery (Figure 1A), the mechanical thresholds of μ^-/-^ mice receiving either vehicle or morphine were significantly lower than baseline mechanical thresholds up to and including day 15 following injection with CFA. Together, these data demonstrate that morphine does not cause anti-hypersensitivity or reinstatement of mechanical hypersensitivity following repeated injections in μ^-/-^ mice.

By contrast, morphine caused anti-hypersensitivity in δ^-/-^ mice significantly elevating mechanical thresholds compared to δ^-/-^ mice receiving vehicle. Mechanical sensitivities were restored back to their sensitivities prior to injection with CFA by day 5 in δ^-/-^ mice receiving morphine, while those receiving saline recovered by day 15. A two-way ANOVA with repeated measures detected a significant effect of time (F8,128 = 25.4; p < 0.0001) and treatment (F1,16 = 7.4; p = 0.015), and a significant interaction (F8,128 = 5.4; p < 0.0001). Pairwise comparisons with the Bonferroni correction revealed that δ^-/-^ receiving morphine had significantly reduced mechanical sensitivities on days 3 – 7 following CFA injection compared to vehicle treated mice (Figure 4B). In contrast to the effect of repeated morphine in WT mice (Figure 3B; Table 2), the capacity of morphine to limit mechanical hypersensitivity in δ^-/-^ mice was sustained up to and including the final measurement on day 15. These data demonstrate that morphine causes anti-hypersensitivity without reinstatement of mechanical hypersensitivity in δ^-/-^ mice despite repeated injections.

### Reinstatement of mechanical hypersensitivity occurs at an accelerated rate in μ^+/-^ mice

Morphine antinociceptive tolerance develops at an accelerated rate and is more profound in μ^+/-^ mice which retain 50% of WT (μ^+/+^) receptor expression (Matthes et al., 1996; Bull et al., 2017a). Therefore, we tested whether reinstatement of mechanical hypersensitivity similarly occurs at an accelerated rate in μ^+/-^ mice by assessing their mechanical thresholds while receiving daily injections of morphine (3 mg/Kg) or vehicle starting from day 2 following injection with CFA (Figure 5).

**Figure 5.**
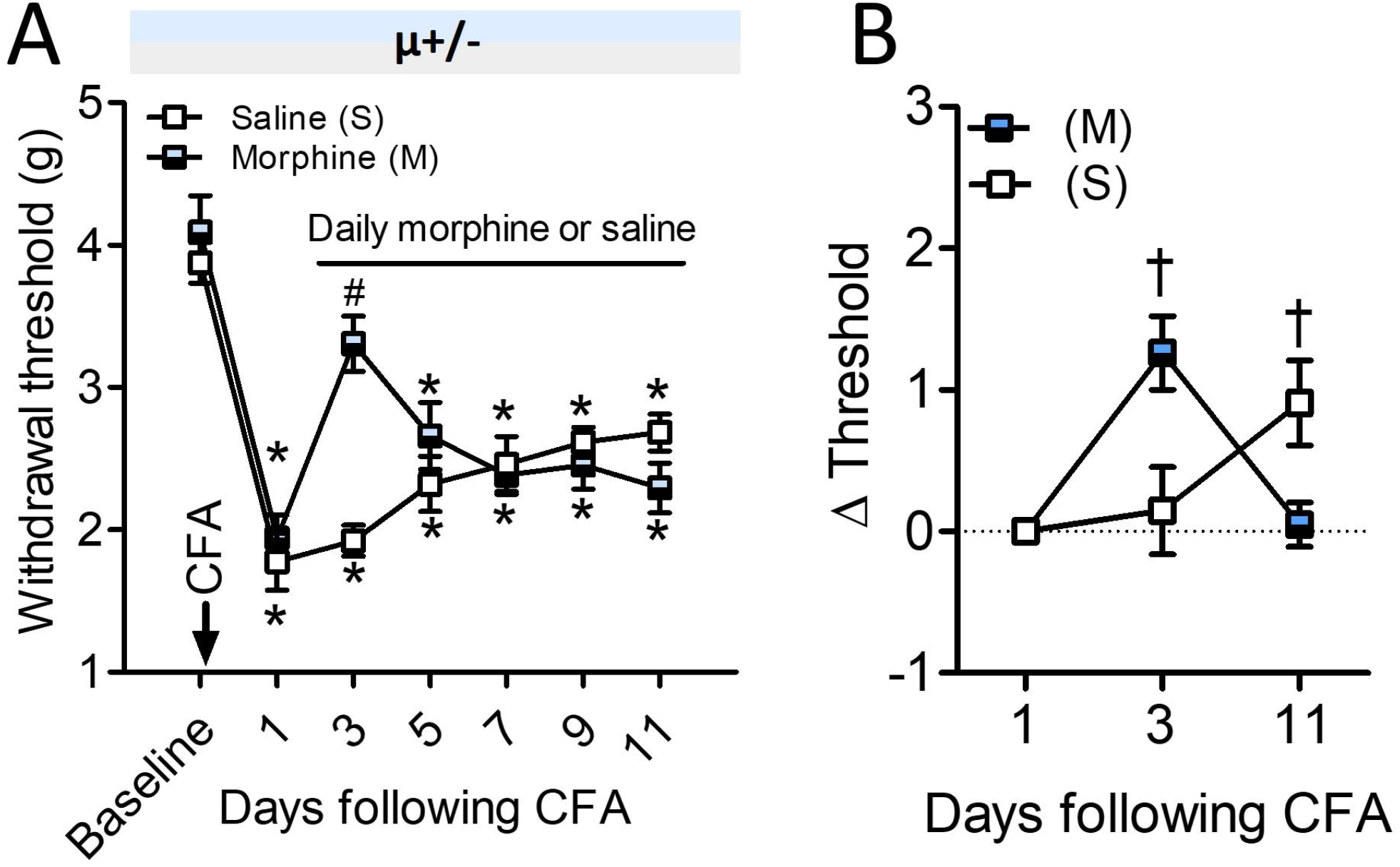
Morphine anti-hypersensitivity and transition to mechanical hypersensitivity in μ^+/-^ mice. (A) Mechanical sensitivities of μ^+/-^ mice receiving once-daily injections with morphine (3 mg/Kg) or an equal volume of saline, starting from day 2 after an injection with 10 μL CFA (1 mg/mL) into one hind paw. Morphine initially caused anti-hypersensitivity on day 3 compared to saline treated mice. However, repeated injections caused hypersensitivity to recur rapidly by day 5. The mechanical thresholds of μ^+/-^ mice receiving saline were significantly reduced from baseline on all days following injection with CFA. (B) Mechanical thresholds expressed relative to day 1 following injection with CFA. A two-way ANOVA with repeated measures revealed that, while morphine limited mechanical hypersensitivity on day 3, complete reinstatement occurred by day 11. By contrast, mechanical hypersensitivity in µ+/- mice receiving vehicle was reduced by day 11 (interaction F2,16 = 14.6, p < 0.001). Data represent mean ± SEM. n = 3 (saline), n = 8 (morphine). *p < 0.05 compared to baseline, ^#^p < 0.05 compared to saline, ^†^ p < 0.05 compared to day 1 sensitivity.

Morphine caused anti-hypersensitivity on day 3 in μ^+/-^ mice elevating mechanical thresholds back to baseline sensitivities whereas saline injections had no effect (Figure 5A). However, mechanical hypersensitivity recurred by day 5 in mice receiving morphine, after receiving just 3 daily injections. Mechanical sensitivities of μ^+/-^ mice on day 3 (after acute injection) and day 11 (after 9 daily injections) were additionally expressed relative to day 1 thresholds to test whether reinstatement of mechanical hypersensitivity is more profound in μ^+/-^ mice following repeated morphine injections (Figure 5B). A two-way ANOVA comparing the change in mechanical sensitivity, from day 1 in µ^+/-^ mice (Δ threshold), detected a significant interaction between time and treatment (F2,16 = 14.6, p < 0.001). Pairwise comparisons with the Bonferroni correction revealed that, in µ^+/-^ mice receiving saline, there was no significant change in mechanical thresholds on day 3 whereas by day 11, mechanical thresholds were significantly elevated compared to day 1. By contrast, while morphine reduced mechanical hypersensitivity on day 3 compared to day 1, no significant difference in thresholds was present by day 11 (Figure 5B).

In comparison, a two-way ANOVA with repeated measures comparing the change in withdrawal thresholds in µ^+/-^ mice to those in WT mice also receiving morphine (3 mg/Kg) revealed that the mechanical thresholds on day 11 in the latter remained significantly higher than day 1 (F2,30 = 5.3, p = 0.01). Furthermore, the mechanical thresholds of WT mice receiving morphine were significantly higher than µ^+/-^ mice, also receiving morphine, on day 11. Together, these data demonstrate that mechanical hypersensitivity is more profoundly reinstated and occurs at an accelerated rate in μ^+/-^ compared to WT mice following repeated injections with morphine.

### Contribution of β-arrestin2 to the modulation of hypersensitivity by morphine

The scaffolding and signalling protein, β-arrestin2, is implicated in morphine tolerance (Bohn et al., 2000). A lack of β-arrestin2 renders mice resistant to tolerance caused by daily administration of 10 mg/Kg morphine (Bohn et al., 2000; Bull et al., 2017a). Therefore, in the experiments that follow we increased the daily dose of morphine and examined its effect on CFA-evoked hypersensitivity. Mice received daily injections with either morphine (10 mg/Kg) or vehicle. Morphine (10 mg/Kg) initially reversed the development of mechanical hypersensitivity in WT mice by day 3, however hypersensitivity recurred on day 9 (Figure 3B; Table 2).

Morphine (10 mg/Kg) also reversed mechanical hypersensitivity evoked by CFA in β-arrestin2^-/-^ mice, again restoring baseline mechanical sensitivities on day 3 (Figure 6). In contrast to β-arrestin2^-/-^ mice that received morphine, mice that received vehicle developed mechanical hypersensitivity that recovered by day 9. A two-way ANOVA with repeated measures confirmed a significant effect of time (F8,112 = 14.7; p < 0.0001) and treatment (F1,14 = 11.3; p = 0.005), as well as detecting a significant interaction (F8,112 = 3.8; p = 0.0005). Pairwise comparisons with the Bonferroni correction revealed that morphine significantly enhanced the mechanical thresholds of β-arrestin2^-/-^ mice compared to vehicle, on days 3 – 7 (Figure 6). Furthermore, the effect of morphine was sustained up to and including the final measurement on day 15 without evidence of developing MIH when compared to vehicle (p = 0.67).

**Figure 6.**
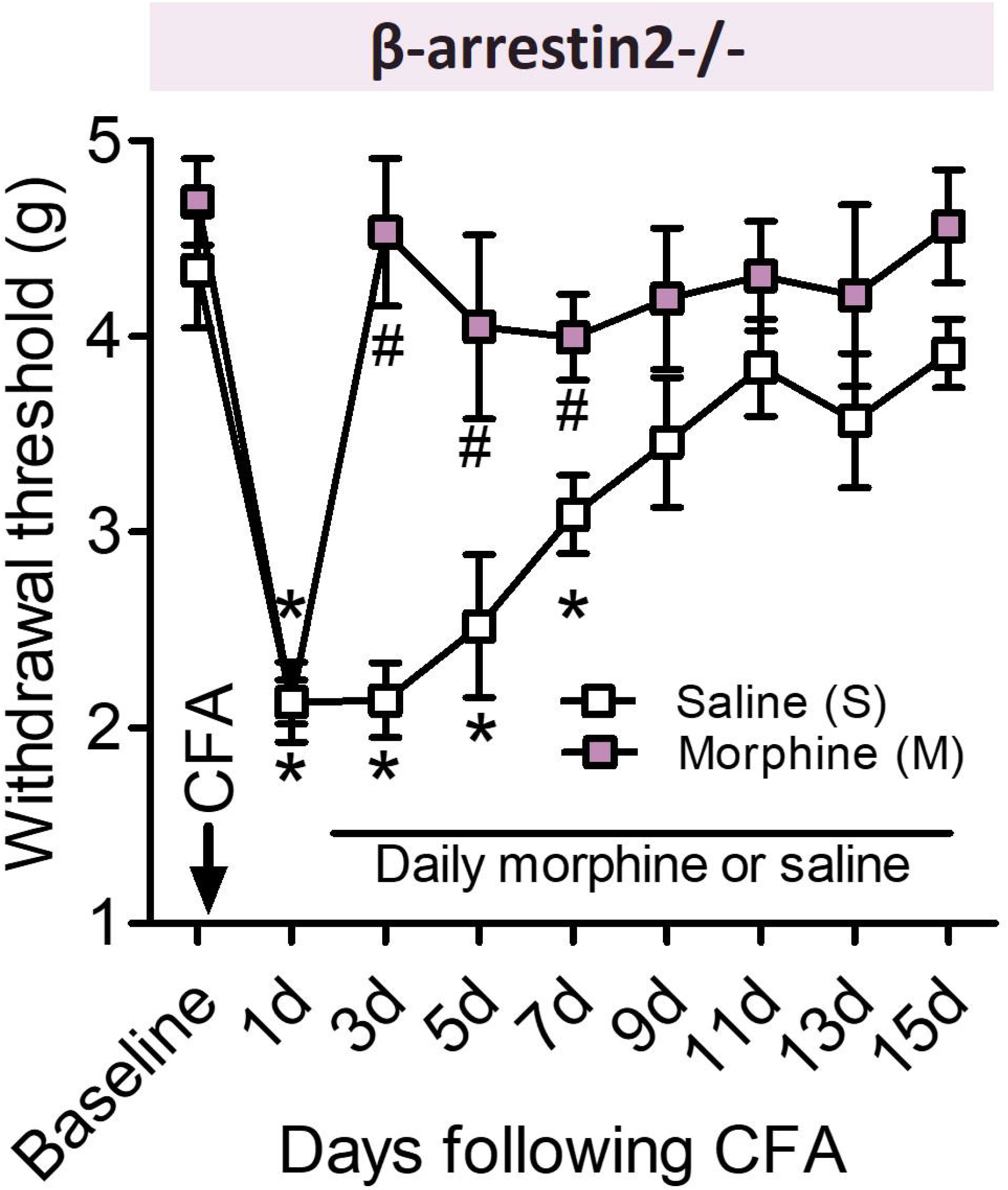
Morphine causes sustained anti-hypersensitivity in β-arrestin2^-/-^ mice. (A) Mechanical sensitivities of β-arrestin2^-/-^ mice receiving once-daily injections with morphine (10 mg/Kg) or an equal volume of saline, starting from day 2 after an injection with 10 μL CFA (1 mg/mL) into one hind paw. A two-way ANOVA with repeated measures revealed that morphine limited hypersensitivity on days 3 – 7 compared to vehicle (treatment: F1,14 = 13.2, p = 0.003; interaction: F2,28 = 41.4, p < 0.001). Furthermore, morphine anti-hypersensitivity was sustained despite repeated injections. (B) Mechanical sensitivities of WT (Figure 3; Table 2) and β-arrestin2-/- mice receiving morphine (10 mg/Kg) expressed relative to baseline. A two-way ANOVA with repeated measures revealed that repeated morphine injections significantly enhanced the mechanical sensitivities of WT mice compared to those of β-arrestin2-/- mice from day 11 (F8,112 = 5.8, p < 0.001). Data represent mean ± SEM. (A) n = 8 (saline), n = 8 (morphine). (B) n = 8 (WT), n = 8 (β-arrestin2-/-). *p < 0.05 compared to baseline, ^#^p < 0.05 compared to saline, ^†^p < 0.05 genotype effect.

By contrast, pairwise comparisons with the Bonferroni correction revealed that the withdrawal thresholds of WT mice receiving morphine (10 mg/Kg; Table 2) were significantly reduced compared to those of β-arrestin2-/- mice (Figure 6) from day 11 (F8,112 = 5.8, p < 0.001). Together, these data demonstrate that mechanical hypersensitivity following repeated morphine injections does not develop in the absence of β-arrestin2.

### c-Src inhibition by dasatinib accelerates recovery from inflammatory hypersensitivity

The non-receptor tyrosine kinase, c-Src, has also been implicated in both inflammatory hypersensitivity and morphine-evoked tolerance (Liu et al., 2008; Xu et al., 2012; Bull et al., 2017a; Chen et al., 2020). Therefore, we determined the consequence of c-Src activation during persistent inflammation using the Src inhibitor, dasatinib (5 mg/Kg), administered intraperitoneally once-daily to WT mice either 2 days after (Figure 7A-C) or 1 day prior (Figure 7D) to injection with CFA. Using the latter approach, we additionally assessed the impact of c-Src inhibition alone on baseline mechanical sensitivities.

**Figure 7.**
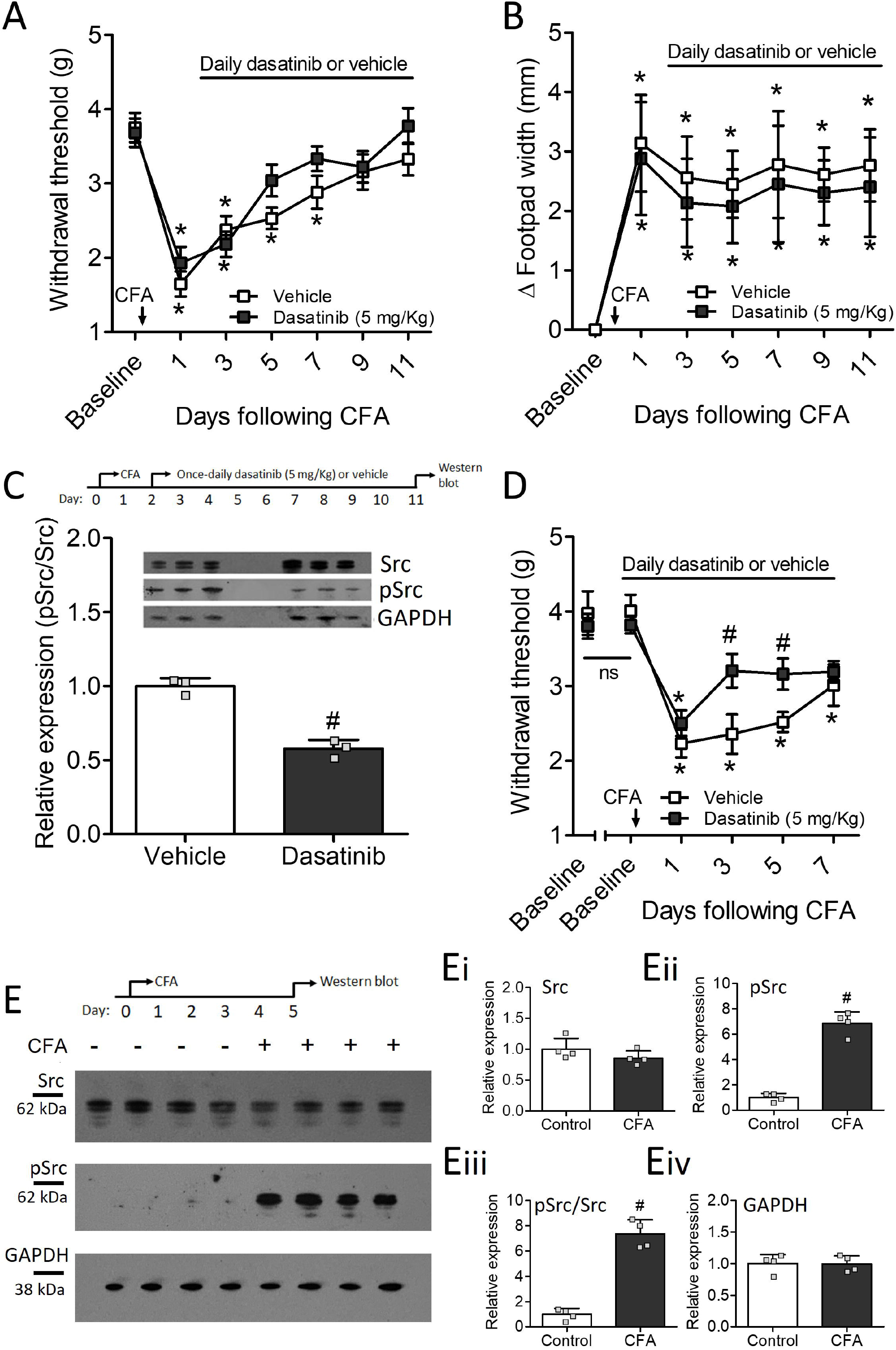
Dasatinib accelerates recovery from mechanical hypersensitivity without affecting baseline hypersensitivity or inflammation. WT mice were injected with 10 μL CFA (1 mg/mL) to test the consequences of c-Src activation on mechanical hypersensitivity during persistent inflammation. (A) Mechanical sensitivities of WT mice receiving once-daily injections with vehicle (saline supplemented with 2% v/v DMSO and 2% v/v Kolliphor EL) or dasatinib (5 mg/Kg dissolved in vehicle). A two-way ANOVA revealed that dasatinib significantly accelerated recovery back to baseline sensitivities by day 5 whereas recovery was delayed until day 9 in mice receiving vehicle. This occurred without changes to footpad inflammation (B) and was accompanied by reduced c-Src phosphorylation within the lumbar spinal cord (C). There was no effect on baseline mechanical sensitivities when dasatinib was administered prior to injection with CFA (D), however, daily injections of dasatinib similarly accelerated recovery back to baseline sensitivity by day 3. (E) Western blot using 10 µg of total protein from spinal cord tissue harvested 5 days after injection with saline (4 left columns) or CFA (4 right columns) revealed that, while c-Src was predominantly in its non-phosphorylated, inactive form, in the spinal cords of mice without CFA (-), c-Src phosphorylation (Y416), and hence, activation, was detected in the spinal cords of mice with CFA (+). See Methods for all details of protein sample preparation, western blotting and detection. Each lane is the tissue from an individual mouse. Data are expressed as mean ± S.E.M (A and D) or mean ± S.D (B, C, E_i-iv_). n (per treatment) = 16 (A), 8 (B), 3 individual mice (C), 10 (D), and 4 individual mice (E). *p < 0.05 compared to baseline, ^#^p < 0.05 treatment effect.

Dasatinib accelerated recovery from mechanical hypersensitivity when administered once-daily following CFA injection (Figure 7A). A two-way ANOVA with repeated measures detected a significant interaction between time and treatment (F6,180 = 27.4; p = 0.04). Pairwise comparisons with the Bonferroni correction revealed that the mechanical thresholds of mice receiving dasatinib were significantly lower than baseline up to but not including day 5, whereas recovery back to baseline mechanical sensitivity was delayed until day 9 in mice receiving vehicle. This occurred without any effect of dasatinib on footpad width (Figure 7B).

We performed western blots on the spinal cords of mice injected with CFA that had received once-daily injections with dasatinib (5 mg/Kg) or vehicle to assess c-Src phosphorylation evoked by CFA in the spinal cord (Figure 7C). Densitometric analysis revealed that total c-Src expression in the spinal cord was similar in mice receiving dasatinib or vehicle. However, the relative expression of active pSrc, identified using the antibody against its Y416 phosphorylated form (Roskoski, 2005), was significantly reduced by dasatinib. Furthermore, the ratio of phosphorylated to non-phosphorylated c-Src, a measure of relative c-Src activation, confirmed that dasatinib reduced c-Src activation in the spinal cord despite persistent inflammation (unpaired student’s T-test, p = 0.0008). These data demonstrate that dasatinib accelerates the recovery from mechanical hypersensitivity that coincides with a reduction in c-Src phosphorylation within the spinal cord.

We next assessed the effect of dasatinib (5 mg/Kg) on baseline mechanical sensitivities when administered prior to injection with CFA (Figure 7D). Baseline thresholds before and after injection with vehicle were: 4.0 ± 0.3 g and 4.0 ± 0.2 g whereas baseline thresholds before and after injection with dasatinib were 3.8 ± 0.2 g and 3.8 ± 0.1 g. A two-way ANOVA with repeated measures comparing the mechanical thresholds before and after mice had received either vehicle or dasatinib revealed no significant effect of time (F1,18 = 0.1; p = 0.92) or treatment (F1,18 = 1.2; p = 0.28). These data demonstrate that dasatinib does not affect mechanical sensitivity in the absence of inflammation. We then injected mice with CFA while continuing to administer once-daily injections with dasatinib or vehicle. We observed a similar trend in the recovery from mechanical hypersensitivity when dasatinib was administered prior to the injection of CFA (Figure 7D) as we did when dasatinib was administered after injection with CFA (Figure 7A). However, recovery back to baseline sensitivity occurred by day 3 in dasatinib pre-treated mice. A two-way ANOVA with repeated measures comparing mechanical sensitivities confirmed a significant effect of treatment (F1,18 = 5.3; p = 0.033). Pairwise comparisons with the Bonferroni correction revealed that dasatinib caused anti-hypersensitivity (compared to vehicle) on day 3 and day 5 following injection with CFA (Figure 7D). Together, these data demonstrate that dasatinib reduces inflammatory hypersensitivity when administered prior to, and after, CFA.

### Complete Freund’s adjuvant evokes c-Src kinase activation in the spinal cord

Since CFA administration into the hind paw reduces Csk mRNA expression in the spinal cord (Figure 2H), and c-Src inhibition by dasatinib accelerated recovery by day 5 (Figure 7A), we tested whether inflammation evoked by CFA causes the activation of c-Src kinase within the spinal cord of WT mice (Figure 7E). Mice received CFA or vehicle (saline) and their lumbar spinal cords were harvested after 5 days. Total c-Src expression, and c-Src phosphorylation at Y^416^, was determined by western blotting. Together, these were used to calculate the ratio of phosphorylated, active c-Src, to total c-Src, a measure of c-Src activation (Figure 7C).

CFA increased c-Src activation in the lumber spinal cord (Figure 7E). Densitometric analysis of the visualised bands revealed that total c-Src expression was similar in the spinal cords of mice that received CFA compared to those that received vehicle (Figure 7E_i_; unpaired T-test, p = 0.22). In contrast, the level of phosphorylated c-Src was virtually undetectable at baseline whereas expression was significantly enhanced following 5 days of inflammation as indicated by the relative expression of pSrc to GAPDH (Figure 7E_ii_; p < 0.0001) and the pSrc/Src ratio (Figure 7E_iii_; p < 0.0001). There was no change in GAPDH expression between vehicle and CFA injected mice confirming equal loading (Figure 7E_iv_; p = 0.96). These data demonstrate that persistent CFA-evoked inflammation activates signalling pathways which cause c-Src activation within the spinal cord.

### Dasatinib prevents reinstatement of mechanical hypersensitivity following morphine injection

Daily administration of dasatinib (5 mg/Kg) attenuates the development of morphine (10 mg/Kg) antinociceptive tolerance assessed using tail withdrawal from hot water (Bull et al., 2017a). We determined whether dasatinib prevents hypersensitivity from recurring following repeated injections with this same dose of morphine in mice with persistent inflammation (Figure 8).

**Figure 8.**
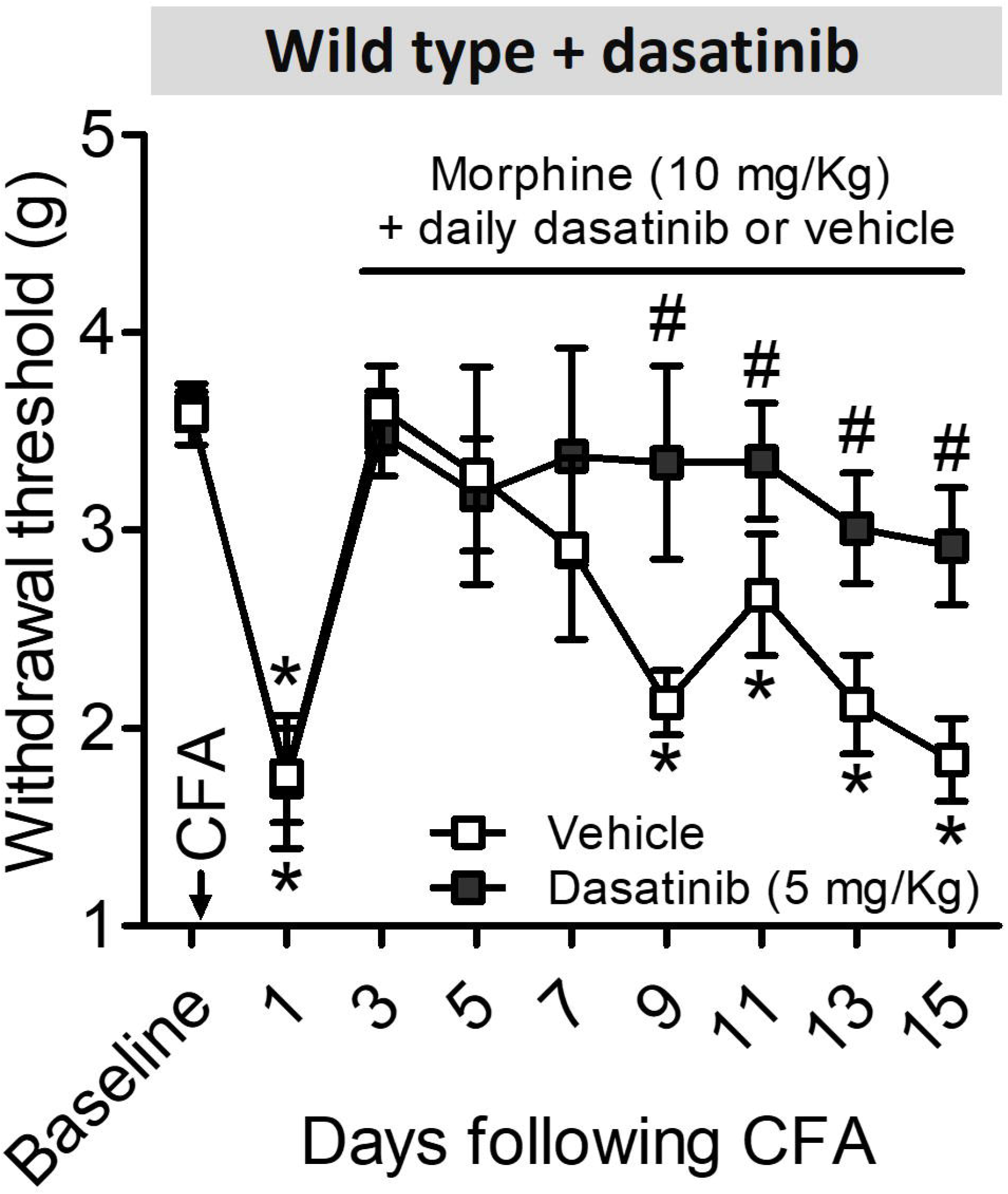
Dasatinib enhances morphine anti-hypersensitivity by preventing the transition to mechanical hypersensitivity. (A) Mechanical sensitivities of WT mice receiving once-daily injections (i.p) of either vehicle (saline supplemented with v/v 2% DMSO and 2% Kolliphor EL) or dasatinib (5 mg/Kg) 30 minutes prior to receiving chronic, once-daily injections of morphine (10 mg/Kg; s.c) starting from day 2 after injection with complete Freund’s adjuvant (CFA). Morphine caused anti-hypersensitivity in all mice on day 3. However, repeated morphine injections caused hypersensitivity to recur in mice receiving prior injection with vehicle while morphine anti-hypersensitivity was sustained in mice receiving dasatinib. Mechanical sensitivities of mice receiving vehicle were significantly enhanced from day 9 onwards compared to mice receiving dasatinib. Data represent the mean ± SEM. n = 8 (vehicle), n = 8 (dasatinib). *p < 0.05 compared to baseline, ^#^p < 0.05 treatment effect.

Morphine (10 mg/Kg) initially reversed the mechanical hypersensitivity evoked by injection with CFA in mice receiving intraperitoneal vehicle 30 minutes prior to morphine. However, once-daily injections of morphine repeated across consecutive days led to the reappearance of mechanical hypersensitivity in control mice, while mechanical hypersensitivity did not reappear in mice receiving intraperitoneal dasatinib (5 mg/Kg) prior to morphine. A two-way ANOVA with repeated measures detected a significant effect (F8,112 = 6.8; p = 0.004). Pairwise comparisons with the Bonferroni correction revealed that the mechanical thresholds of mice receiving vehicle were significantly reduced compared to baseline, and to the mechanical thresholds of mice receiving dasatinib, from day 9 - 15. In contrast, mice receiving dasatinib had significantly reduced mechanical thresholds from baseline on day 1 following injection with CFA (Figure 8). These data demonstrate that prior administration of the Src inhibitor, dasatinib, prevents reinstatement of mechanical hypersensitivity observed following repeated morphine injections.

## Discussion

We identified roles for several components of the opioid pathway already implicated in morphine tolerance, in inflammatory hypersensitivity, the morphine antihypersensitivity and MIH. Mice lacking expression of µ or δ receptors, or WT mice treated with the Src inhibitor, dasatinib, had similar CFA-evoked inflammation and initial hypersensitivity compared to control WT mice although spontaneous recovery was altered. Spontaneous recovery from mechanical hypersensitivity in WT mice (in the absence of morphine) was complete by day 7 whereas recovery was abolished in µ-/- mice and delayed until day 13 in δ-/- mice. In contrast, WT mice treated with dasatinib had accelerated recovery when compared to vehicle.

Targeting components of the opioid pathway led to altered morphine responses following repeated injections. Daily morphine initially limited CFA-evoked hypersensitivity in WT mice, although mechanical hypersensitivity was reinstated after repeated injections, revealing MIH. Neither of these effects were observed in µ-/- mice confirming the requirement for µ opioid receptors in MIH (Matthes et al., 1996; Corder et al., 2017; Roeckel et al., 2017). By contrast, morphine caused antihypersensitivity in µ^+/-^ mice with an accelerated transition to MIH. Fast MIH likely results from the rapid development of morphine antinociceptive tolerance in µ^+/-^ mice (Sora et al., 2001; Bull et al., 2017a). Conversely, conditions that abolish morphine antinociceptive tolerance, i.e., δ^-/-^ and β-arrestin2^-/-^, or WT mice administered the c-Src inhibitor, dasatinib (Zhu et al., 1999; Bohn et al., 2000; Bohn et al., 2002; Nitsche et al., 2002; Bull et al., 2017a), prevented MIH. Taken together, these observations are consistent with the hypothesis that, in this model of inflammatory hypersensitivity, MIH occurs through similar mechanisms to those previously established for morphine tolerance. It is likely that morphine tolerance reduces the opioid receptor mediated recovery from CFA-evoked inflammatory hypersensitivity leading to MIH.

Spontaneous recovery from mechanical hypersensitivity in WT mice on day 7 was accompanied in the spinal cord by changes in the levels of transcripts encoding opioid peptide precursors, opioid receptors, β-arrestin2 and C-terminal Src kinase. Proopiomelanocortin and proenkephalin mRNAs decreased after 7 days of CFA-evoked inflammation, while prodynorphin mRNA levels increased. These changes in opioid peptide precursors are consistent with previous reports (Luo et al., 2008; Obara et al., 2009). However, the importance of altered proopiomelanocortin and proenkephalin expression is unclear given that recovery from CFA-evoked hypersensitivity is unaltered in *Pomc*^*-/-*^ and *Penk*^*-/-*^ mice (Walwyn et al., 2016). These findings suggest that peptides derived from proopiomelanocortin and proenkephalin contribute little to spontaneous recovery. In contrast, *Pdyn*^*-* /-^ mice exhibit accelerated recovery from CFA-evoked hypersensitivity suggesting that prodynorphin-derived peptides are pronociceptive (Walwyn et al., 2016). Therefore, enhanced spinal prodynorphin following CFA may oppose spontaneous recovery from CFA-evoked hypersensitivity. Indeed, CFA-evoked hypersensitivity is reduced following intrathecal injection of anti-dynorphin antiserum in rats (Luo et al., 2008). It is possible that accelerated recovery also reflects enhanced basal nociception in *Pdyn*^*-/-*^ mice indicating that prodynorphin derived peptides have tonic antinociceptive effects (Gendron et al., 2007; Walwyn et al., 2016). However, studies using this mouse model identify no change in basal mechanical and thermal sensitivities (Maldonaldo et al., 2018). Furthermore, most studies of persistent nociceptive stimuli, including CFA-evoked inflammation, demonstrate hypersensitivity associated with elevated dynorphin (Podvin et al., 2016). These effects appear to be independent of opioid receptor function and are mediated through activation of either NMDA or bradykinin receptors.

Mechanical hypersensitivity in WT and transgenic mice lacking endogenous opioid peptides that have recovered from CFA-evoked hypersensitivity is reinstated by the µ opioid receptor inverse agonist, naltrexone, but not by a neutral antagonist (Corder et al., 2013; Walwyn et al., 2016). This implies that recovery of mechanical sensitivity is primarily mediated by constitutively active µ opioid receptors and therefore does not require opioid peptide agonists. Consistent with a requirement for µ opioid receptors, in our study mechanical hypersensitivity failed to recover in µ^-/-^ mice following CFA. Interestingly, although there was no change in μ opioid receptor mRNA expression 7 days after CFA administration to WT mice, the level of δ opioid receptor mRNA increased whereas κ receptor, β-arrestin2 and C-terminal Src kinase mRNAs decreased. While it is important to note that changes in mRNA expression measured using qPCR may not translate to altered protein expression within the spinal cord (Ji et al., 1995; Obara et al., 2009; Corder et al., 2013), modified expression of δ receptors, κ receptors, β-arrestin2 and C-terminal Src kinase may contribute to spontaneous recovery from mechanical hypersensitivity and influence the morphine response in WT mice. Indeed, abolition of MIH in δ^-/-^, β-arrestin2^-/-^, and WT mice treated with daily dasatinib confirms roles for these components of the opioid signalling system.

Restoration to basal mechanical thresholds following CFA was delayed in δ^-/-^ compared to WT mice in agreement with previous evidence (Gaveriaux-Ruff et al., 2008; Pradhan et al., 2013). This is consistent with enhanced expression of δ receptors within the spinal cord contributing to recovery from mechanical hypersensitivity. Cahill and colleagues also reported an upregulation of δ receptors in the spinal cord (Cahill et al., 2003), which coincides with enhanced efficacy of SNC80, a δ receptor agonist, in attenuating mechanical and thermal hypersensitivity in mice following CFA (Gaveriaux-Ruff et al., 2008; Gaveriaux-Ruff et al., 2011).

The observation of delayed spontaneous recovery from CFA-evoked hypersensitivity in µ^+/-^ and δ^-/-^ compared to WT mice implies that altered surface expression of these receptors participates in the restoration of mechanical sensitivity. Since µ and δ receptors may associate to form heteromers (Gomes et al., 2000; Manglik et al., 2012), and this occurs in mouse spinal cord (Gomes et al., 2011; He et al., 2011), it is possible that δ receptor expression, induced following CFA-evoked hypersensitivity, may increase the proportion of µ-δ heteromers and this may contribute to recovery to basal sensitivity. Interestingly, CFA does not promote changes in surface δ receptor expression in µ^-/-^ mice (Morinville et al., 2004). This implies that µ and δ receptors interact, either physically by the formation of heteromers or through signaling crosstalk in the spinal cord during CFA-evoked inflammation (Fujita et al., 2015). Importantly, when expressed together with δ receptors, µ receptors exhibit agonist-independent recruitment of β-arrestin2 (Rozenfeld et al., 2007; Bull et al., 2017b) and this may play a role in priming morphine tolerance and MIH. Agonist independent recruitment of β-arrestin2 to µ-δ heteromers during persistent inflammation may additionally enhance hypersensitivity by sequestering β-arrestin2 from TRPV1 receptors (Rowan et al., 2014a; Rowan et al., 2014b).

In contrast to µ^-/-^ and δ^-/-^ mice which exhibited diminished spontaneous recovery, dasatinib caused accelerated recovery from hypersensitivity in WT mice without affecting acute mechanical nociception or initial hypersensitivity. This is consistent with studies of the impact of Src inhibition on inflammatory hypersensitivity in rats (Guo et al., 2002; Liu et al., 2008; Xu et al., 2012; Chen et al., 2020). The observed increase in c-Src phosphorylation in the spinal cord following CFA supports previous studies linking inflammation to c-Src activation (Guo et al., 2002; Liu et al., 2008; Xu et al., 2012). Our qPCR data also revealed that the expression of C-terminal Src kinase mRNA is reduced in the spinal cord after CFA. C-terminal Src kinase-mediated phosphorylation diminishes the activity of c-Src kinase (Harvey et al., 1989; Okada, 2013). Therefore, limited C-terminal Src kinase expression within the spinal cord would be expected to enhance c-Src phosphorylation, and hence, activity. This implies that inflammation enhances the activation of c-Src thereby facilitating hypersensitivity. In the spinal cord, activated c-Src phosphorylates the ND2 subunit of the NMDA receptor increasing its surface expression and function, which may promote hypersensitivity (Liu et al., 2008).

Opioid-evoked recruitment of β-arrestin2 to μ receptors is associated with c-Src activation in mouse dorsal root ganglion neurones (Walwyn et al., 2007). If CFA-evoked inflammation causes an upregulation of µ-δ heteromers then their constitutive recruitment of β-arrestin2 may contribute to neuronal c-Src activation (Rozenfeld and Devi, 2007; Bull et al., 2017a). Furthermore, c-Src activation of GRK2 is involved in agonist-evoked β-arrestin2 recruitment to δ receptors resulting in their desensitisation and internalisation (Audet et al., 2005; Archer-Lahlou et al., 2009; Hong et al., 2009). Approaches that diminish c-Src activity and/or expression abolish desensitisation of δ receptors while enhancing their membrane recovery (Archer-Lahlou et al., 2009). Therefore, c-Src inhibition using dasatinib may hasten recovery from CFA-evoked hypersensitivity by regulating δ receptor function. Irrespective of the mechanisms involved, our findings support the idea that c-Src inhibitors may be effective in treating persistent inflammatory pain (Ge et al., 2020).

Taken together with previous studies (Corder et al., 2013; Walwyn et al., 2016) our findings support the idea that CFA-evoked hypersensitivity causes a compensatory up-regulation of tonic opioid signalling restoring basal mechanical sensitivity. We hypothesise that enhanced opioid signalling, mediated by µ and δ opioid receptors, leads to recruitment of β-arrestin2 and activation of c-Src. This primes the system to morphine evoked tolerance and MIH. If similar mechanisms are at play during inflammatory hypersensitivity in humans, it may be advantageous to minimise exposure to opioid agonists that stimulate recruitment of β-arrestin2.

There has been a concerted effort to develop µ agonists biased against β-arrestin2 recruitment in favour of G protein stimulation leading to the recent approval of oliceridine by the US Food and Drug Administration (Colvin et al., 2019; Ammar et al., 2022). Our findings suggest that the development of such drugs may be beneficial to minimise opioid induced hyperalgesia caused by diminished opioid signalling. However, recent evidence suggests that weak recruitment of β-arrestin2 to the μ receptor is a consequence of oliceridine’s partial efficacy rather than signalling bias and this may not be sufficient to prevent tolerance and other detrimental effects (Gillis et al., 2020; Singleton et al., 2021).

An alternative approach to improve opioid analgesia is inhibition of targets that may interact with β-arrestin2 such as c-Src or NMDA receptors (Chen et al., 2016; Colvin et al., 2019). Inhibitors of c-Src and NMDA receptors may also be analgesic even in the absence of opioid agonists. While not explored in this study, there are several preclinical studies that implicate NMDA receptors in opioid tolerance, hypersensitivity and neuropathic pain (Mao et al., 1995; Liu et al., 2008; Chen et al., 2016; Colvin et al., 2019; Chen et al., 2020). Indeed, the clinically available antagonist and dissociative anaesthetic, ketamine, is often used during the perioperative period to reduce pain and diminish the requirement for opioids (Colvin et al., 2019; Culp et al., 2021).

Taken together with previous studies, our findings suggest that c-Src inhibitors, such as dasatinib, may be effective in treating persistent pain either alone, or, as adjuncts to opioid analgesics to prolong their analgesic effects by diminishing opioid-induced tolerance and hypersensitivity.

## Acknowledgements

We thank the staff of the University of Dundee MSRU for assistance with animal experiments and husbandry. SS and TGH are members of the Advanced Pain Discovery Platform.

